# Interpretable deep generative ensemble learning for single-cell omics with Hydra

**DOI:** 10.1101/2025.08.15.670517

**Authors:** Manoj M Wagle, Chunlei Liu, Zunpeng Liu, Yongheng Wang, Manolis Kellis, Ellis Patrick, Pengyi Yang

## Abstract

Single-cell omics enable the dissection of cellular heterogeneity, yet the high dimensionality, inherent noise, and sparsity present significant challenges. These challenges are amplified for rare cell populations, which are often difficult to annotate reliably but can be central to development and disease. As single-cell assays increasingly capture multiple molecular layers, the integrative analysis of such multimodal data further increases complexity. Here, we propose Hydra, a deep generative framework based on an ensemble of variational autoencoders for effective learning of unimodal and multimodal single-cell omics data. Hydra implements interpretable modules for capturing cell type-specific molecular signatures. The ensemble of such interpretable modules enables reproducible feature selection and robust cell type annotation, with particular effectiveness for rare populations. We benchmarked Hydra on a repertoire of 21 datasets, including unimodal and multimodal single-cell omics data. Our results demonstrate that Hydra offers comparable to superior performance to several state-of-the-art methods. Finally, we highlight the utility of Hydra in robustly annotating brain cellular subtypes and preserving disease-relevant signatures using our previously published dataset that profiles Alzheimer’s disease.

## Introduction

The establishment of single-cell transcriptomics has achieved remarkable success in revealing cell type-specific molecular variations by enabling the profiling of each individual cell (1–3). More recently, there has been a transition toward single-cell multiomics which allows for simultaneous probing of other modalities such as chromatin accessibility and surface proteins from the same cell in a single experiment (4–7). Such “paired data” enables a more comprehensive understanding of cellular function by capturing different layers of biological information, such as epigenetic and signalling regulation. Despite these advances, analyzing single-cell omics data presents its own set of challenges. Single-cell data is often sparse and noisy, with each cell characterized by thousands of features. This high dimensionality complicates the identification of key molecular signatures that underpin the biological variations and hinders downstream analyses such as cell type identification. Consequently, feature selection has become a useful step to identify cell-type specific signatures and improve the signal-to-noise ratio in downstream analyses (8, 9).

Accurate identification of cell types therefore requires the selected features to be reproducible and generalizable across studies and technologies. This is particularly important for supervised cell type prediction frameworks that utilize the selected features from reference datasets to identify cell types in new single-cell studies. A number of such supervised tools have been developed, including statistical and classical machine learning-based approaches, and more recently, deep learning models have gained increasing popularity in this domain due to their ability to handle the complexities of single-cell data (10–12). Nevertheless, three key challenges remain. First, most existing tools are developed specifically for single-cell transcriptomic data and do not account for other single-cell omics modalities, limiting their applicability in multi-omic contexts. Second, existing methods often fail to detect smaller cell populations and are biased toward major cell types, which hinders their ability to recover rare cell populations that may nevertheless play critical roles in development and disease. Finally, deep learning-based approaches often lack interpretability, which makes it difficult to understand their predictions (13, 14).

To bridge this gap, we introduce Hydra, a deep generative framework designed to effectively handle unimodal and multimodal single-cell omics data that profile gene expression, chromatin accessibility, and surface protein abundance. A key strength of Hydra is its ability to reliably recover and annotate rare cell populations, where class imbalance and weak marker signals pose challenges for existing methods. In particular, the feature ranking module of Hydra is based on training an ensemble of variational autoencoders (VAEs) each paired with a classification head for gradient descent–based feature attribution. This ensemble deep learning approach (15) provides interpretable and cell type–specific markers that highlight the key molecular features distinguishing each population (13), and is especially effective for rare cell types where existing methods often fail to recover robust markers.

By learning the probability distributions of the data via VAEs, Hydra can simulate new samples for smaller cell type populations. This enables data augmentation for learning from balanced datasets and thus addresses the issue of class imbalance. The molecular features identified by the feature ranking module of Hydra are utilized by its cell type mapping module, which again employs ensemble learning using simple neural networks to automatically predict cell types in query datasets. The architecture of Hydra is adaptable for both unimodal and paired multimodal single-cell omics data. Using 21 datasets generated from various tissue types and diverse single-cell omics technologies, we demonstrate that Hydra improves feature selection reproducibility and outperforms several popular tools, especially on rare cell type annotation. Together, these results highlight the utility and flexibility of Hydra for feature selection and cell type annotation of diverse unimodal and multimodal single-cell omics data.

## Results

### Overview of Hydra framework

We developed Hydra, an interpretable ensemble deep learning framework for jointly identifying cell type-specific markers and mapping cell types in unimodal data, such as single-cell transcriptomes, as well as multimodal data such as those that jointly measure transcriptomes, chromatin accessibility, and protein expression in individual cells. Specifically, Hydra extends on the variational autoencoder (VAE) architecture utilizing a multi-task training procedure (16) (Fig. 1a), and contains two modules (i) an ensemble feature ranking module (Fig. 1b) and (ii) an ensemble cell type annotation module (Fig. 1c). First, the VAE architecture is jointly trained with an ensemble of classification heads (Fig. 1a). During this phase, we apply a dynamic sampling strategy to address cell type imbalances commonly encountered in single-cell data by using the VAE to generate synthetic cells for minor cell types and random down-sample major cell type populations (Methods). The model is optimized by a loss function that takes into account the reconstruction loss from the decoder and the loss from the classification heads. Hydra also minimizes overfitting on the training data by adopting an early stopping approach. The balanced datasets from this initial training phase are used subsequently in the feature ranking and cell type annotation modules.

**Fig. 1.**
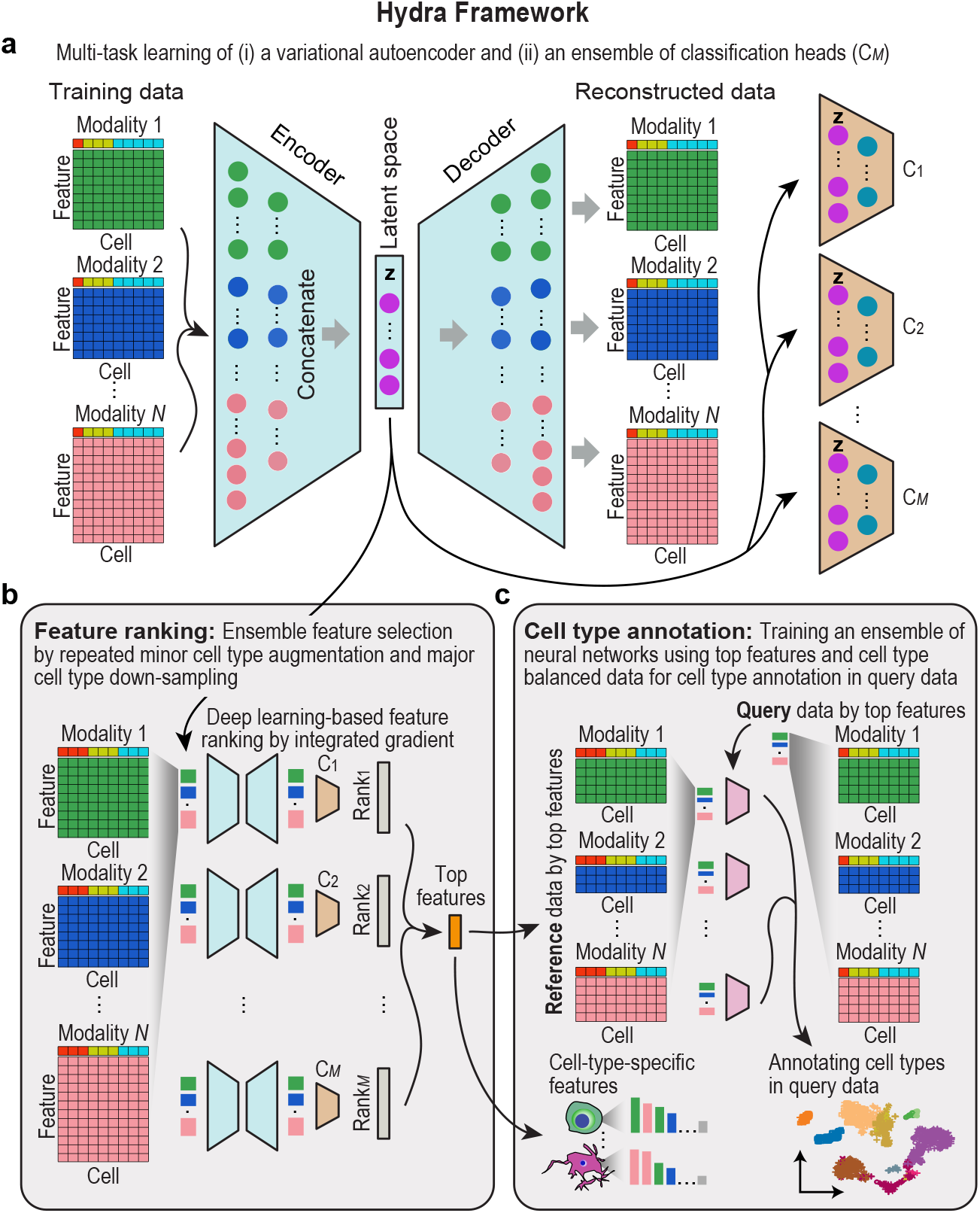
Overview of the Hydra framework for ensemble feature ranking and automated cell type annotation. (a) Graphical illustration of key architectural components in the Hydra framework, including the variational autoencoder (VAE) and the ensemble of classification heads. (b) Schematics of the feature ranking module of the Hydra framework. Data augmentation and down-sampling are used to balance minor and major cell types. Multiple cell type balanced datasets each paired with a classification head is used for feature ranking and the consensus is taken to derive top features. (c) Schematics of the cell type annotation module of the Hydra framework. An ensemble of simple neural networks is trained on dimension reduced and cell type balanced datasets generated from the feature ranking module. These trained neural networks are then employed to predict cell types in any provided query datasets.

In the feature ranking module, we initialize models by inheriting weights from the original trained model and specific classification heads and fine-tune each model using each corresponding balanced dataset (Fig. 1b). Each fine-tuned model is used for feature ranking by employing the post-hoc feature attribution approach of Integrated Gradients (17). This approach accounts for interpretability and obtains cell type-specific ranking of features, considering the biological direction of molecular changes. Finally, feature rankings from individual models are combined to derive consensus top features. In the cell type annotation module, an ensemble of simple neural network classifiers is trained each on a balanced dataset filtered by top features selected in the feature ranking module (Fig. 1c). After training, the predictions of cells to cell types for a given query dataset are averaged across individual classifiers to obtain the final cell type.

### Ensemble deep learning improves feature stability of Hydra

Reliable identification of cell type markers requires feature importance estimates to be consistent across dataset perturbations. To assess whether the ensemble modeling approach implemented in Hydra is useful for improving the stability of feature selection results, we evaluated ensemble sizes of 10, 25, 50, and 100 and compared their stability to that of a single model where no ensemble is not used. Specifically, we performed stratified subsampling based on original cell type proportions using two independent single-cell transcriptomic (scRNA-seq) lung datasets (18, 19) to create different variants of the original datasets. We then performed feature selection using the above models and computed pairwise Pearson correlation coefficients of feature importance from these variants for each dataset. These correlations were used to measure stability of features selected. We found that the ensemble implementation of Hydra demonstrated a higher feature selection stability compared to the single model. Specifically, we did not observe any significant improvement in stability beyond an ensemble size of 25 (Fig. 2a, Supplementary 1a-b). Therefore, we selected an ensemble size of 25 as the default configuration for all subsequent analyses (we refer to this configuration simply as “Hydra” throughout the manuscript).

**Fig. 2.**
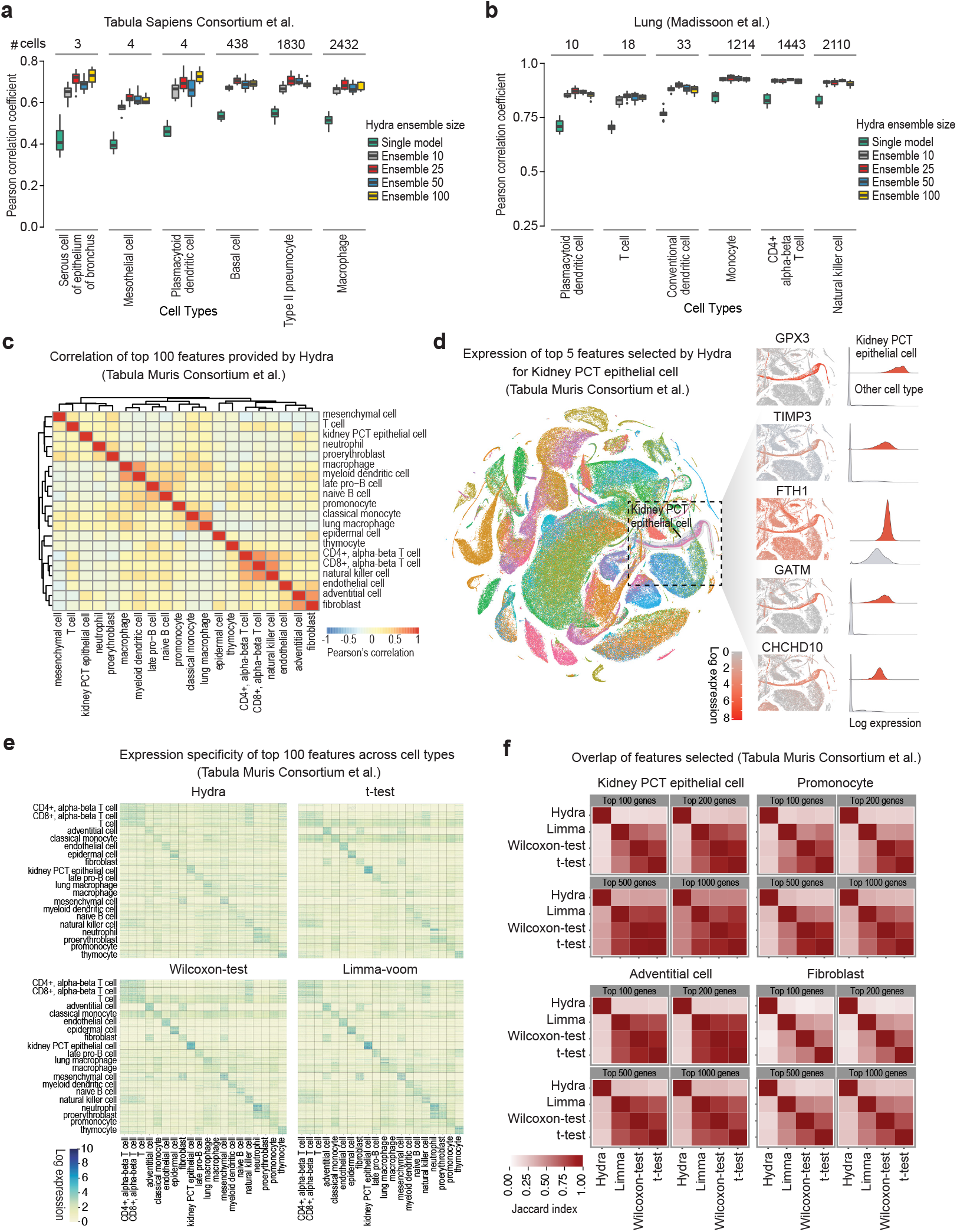
Evaluation of Hydra feature stability, cell type specificity, and correlation across different methods. (a, b) Feature selection stability of Hydra quantified using Pearson’s correlation coefficient using different ensemble sizes (n = 1, 10, 25, 50, 100) on (a) Tabula Sapiens and (b) Lung datasets. The three smallest and three largest cell types from each dataset are shown, arranged in the order of increasing sample count from left to right. (c) Hierarchical clustering of the cell types based on the top 100 features selected by Hydra for 20 cell types using the subsampled dataset derived from the Tabula Muris dataset. The dataset comprises ten major cell types and ten minor cell types, with an imbalance ratio of major to minor cells of 100:2. (d) t-SNE projection of the Tabula Muris dataset highlighting the log-normalized expression of the top five genes selected by Hydra for the kidney PCT epithelial cell type. Additionally, ridge line plots show the specific log-normalized expression of each top gene in kidney PCT epithelial cells compared to all other cell types. (e) Heatmap of log-normalized expression illustrates the cell type specificity of the top 100 genes selected by different methods using the subsampled datasets derived from the Mouse Cell Atlas. The dataset comprises ten major cell types and ten minor cell types, with an imbalance ratio of major to minor cells of 100:2. Columns correspond to cells, and rows represent individual genes specific to their respective cell types. (f) The overlap of top genes (n = 100, 2400, 500, 1000) selected by different methods is quantified using the Jaccard index for four cell types, including kidney PCT epithelial cell, promonocyte, adventitial cell, and fibroblast.

### Capturing cell type-specific markers

We created a subsampled dataset from the scRNA-seq Mouse Cell Atlas (20) consisting of 20 randomly selected cell types with a major-to-minor cell type sample imbalance ratio of 100:2. We then performed feature selection on this dataset using Hydra and three other statistical methods - Welch’s t-test, Wilcoxon rank-sum test, and Limma-Voom (Methods). For each method, we computed Pearson correlations for the top 100 features from each cell type and performed hierarchical clustering. We found that the features selected by Hydra grouped biologically similar cell types together and exhibited strong correlations (Fig. 2c, Supplementary 2a-c). However, the other three methods, including t-test, Wilcoxon, and Limma-voom, failed to group naive B-cell and late pro B-cell together. As another example, Limma-voom and Wilcoxon failed to group classical monocytes and pro-monocytes together.

Next, to demonstrate Hydra’s ability to select cell type-specific features, we performed feature selection on the same subsampled dataset created earlier. Initially, we focused on one cell type - kidney proximal convoluted tubule (PCT) epithelial cells. We selected the top five genes (GPX3, TIMP3, FTH1, GATM, and CHCHD10) identified by Hydra for this cell type and visualized the expression of these genes in the original Mouse Cell Atlas. We found that all five genes were highly expressed in kidney PCT epithelial cells compared to all other cell types, indicating cell type specificity (Fig. 2d). Similarly, we selected and evaluated the expression of the top 100 genes for all 20 cell types from the subsampled dataset. We observed that Hydra displayed a clear diagonal pattern of expression in the heatmap, indicating strong specificity with minimal expression in other cell types (Fig. 2e). While the t-test, Wilcoxon-test, and Limma-Voom also demonstrated similar patterns, the top genes selected by these methods for a few cell types also exhibited expression in non-diagonal blocks. Finally, we assessed the overlap of the top genes selected by different methods for four example cell types - Kidney PCT epithelial cell, Promonocyte, Adventitial cell, and Fibroblast. Specifically, we computed the Jaccard index for the overlap of the top 100, top 200, top 500, and top 1000 genes. Overall, we found that Hydra selected more distinct genes and had a lower overlap with the top genes selected by other methods. In contrast, all statistical methods selected similar sets of genes, as indicated by a higher Jaccard index (Fig. 2f).

### Accurate prediction of cell types in single-cell transcriptomic data across multiple tissues

We next evaluated the performance of Hydra in identifying cell types across a variety of scRNA-seq datasets. We utilized a total of 13 scRNA-seq datasets of varying sizes and numbers of cell types across different platforms and tissues (Table 1). We compared Hydra against four other methods commonly used for predicting cell types in scRNA-seq data - SingleCellNet (21), scClassify (22), scPred (23) and scVI (24). To this end, we adopted intra-dataset and inter-dataset analyses to comprehensively assess the methods. To account for noise, we filtered out genes expressed in less than 1% of the cells in each dataset. All methods were implemented with default parameters following the author’s official documentation. For intra-dataset cell type prediction, we conducted five times repeated random subsampling validation using two datasets - Prostate Urethra (24 cell types, query size=49,000 cells) (25) and Colon (24 cell types, query size=33,000 cells) (26). We found that Hydra outperformed all other methods in predicting most cell types in both datasets, including minor cell types (Fig. 3a-c, Supplementary 3a-b). Notably, all other methods perform reasonably well in identifying larger cell type populations, but their performance decreased for the minor cell types. For example, the Plasmacytoid dendritic cell is the smallest cell type in both datasets. Hydra and scClassify identify this cell type with high accuracy in the colon dataset. However, in the Prostate Urethra dataset, scClassify exhibited significantly lower prediction accuracy for this cell type compared to Hydra.

**Table 1.**
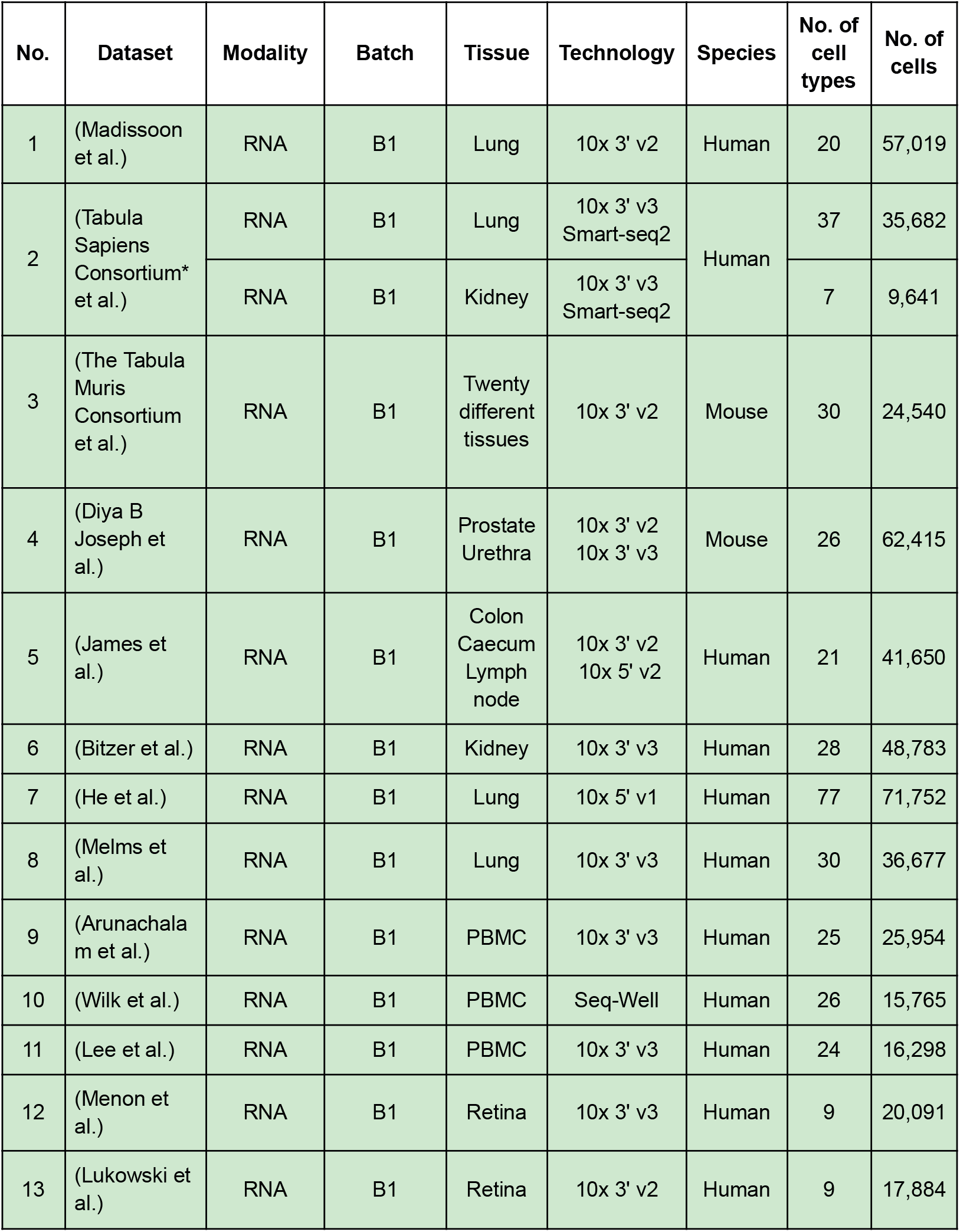
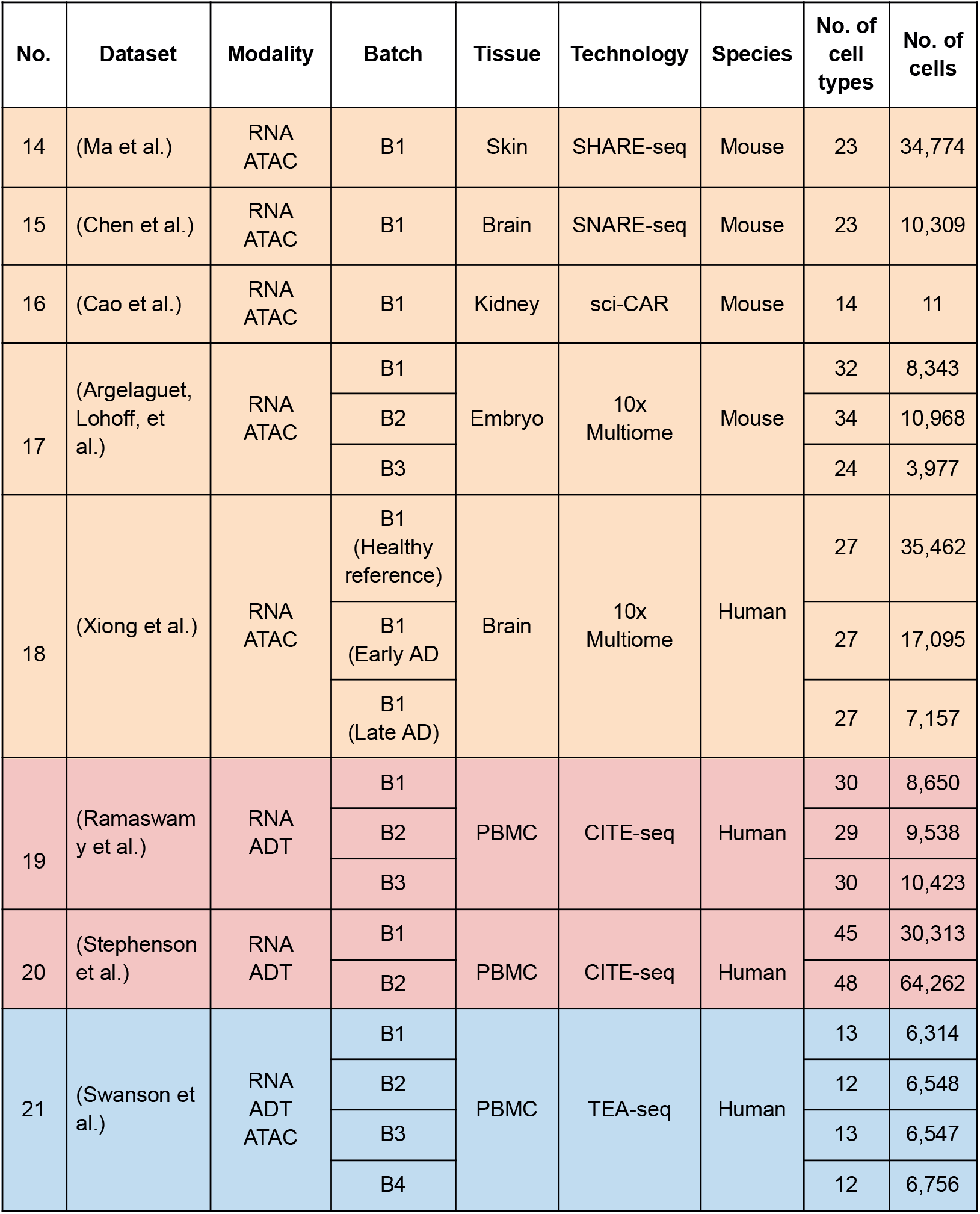
Summary of all datasets used in this study.

**Fig. 3.**
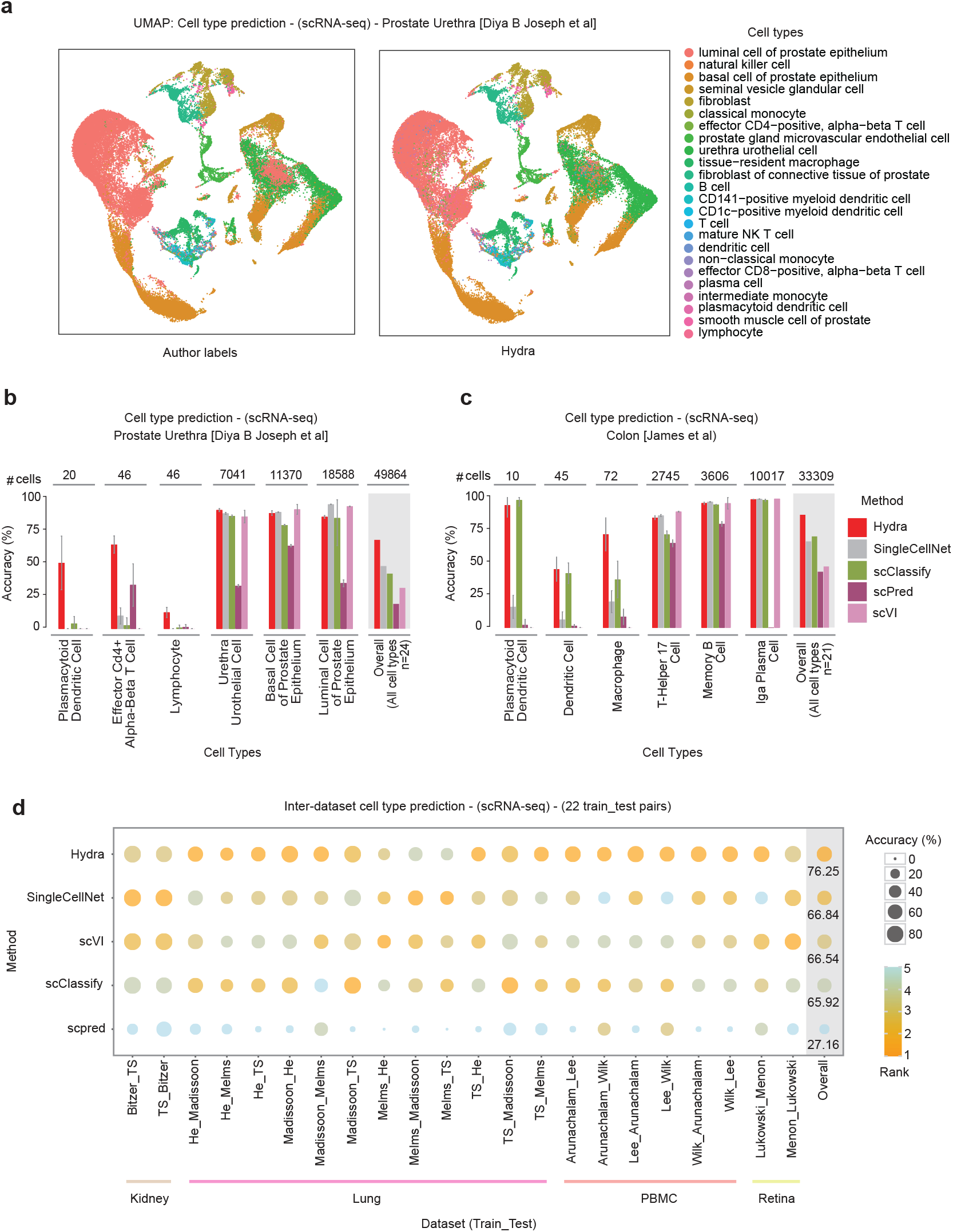
Cell type prediction performance of Hydra against state-of-the-art methods using single-cell transcriptomic datasets. (a) Uniform Manifold Approximation and Projection (UMAP) plots for the single-cell transcriptomic prostate dataset, comprising 24 cell types and approximately 49,000 cells. The left panel shows the original labels as provided by the authors of the study, while the right panel shows the cell types predicted by Hydra. (b) Bar plot with error bars illustrating the five-time repeated random subsampling intra-dataset cell type prediction performance of various methods using the single-cell transcriptomic prostate dataset. Accuracy is computed for each cell type. The three smallest and three largest cell types are displayed, arranged in order of increasing sample count from left to right, followed by the overall performance across all cell types in the dataset (highlighted with a gray background). (c) Bar plot with error bars representing the five-time repeated random subsampling intra-dataset cell type prediction performance of different methods using a single-cell transcriptomic colon dataset comprising approximately 33,000 cells. Accuracy is computed for each cell type. The three smallest and three largest cell types are displayed, arranged in order of increasing sample count from left to right, followed by the overall performance across all 261 cell types in the dataset (highlighted with a gray background). (d) Bubble plot illustrating the inter-dataset cell type prediction performance of various methods across four tissues (Kidney, Lung, PBMC, and Retina) and 22 train-test pairs. Methods are ranked based on their overall mean accuracy across all datasets.

In real-world scenarios, data often come from different sources, which introduces variability and evaluations based on splitting a dataset into train and test sets may not fully capture the generalizability of the methods. To address this, we extended our analyses to inter-dataset cell type prediction tasks where we train and test on independent datasets (e.g., technologies, studies). In total, we used 10 different scRNA-seq datasets for inter-dataset benchmarking across 22 train-test pairs and 4 different tissues, including Kidney, Lung, PBMC and Retina (Table 1). The classification results from this revealed that Hydra consistently achieved a higher prediction performance across most of the datasets (Fig. 3d) compared to other methods. Taken together, the results suggest that Hydra outperforms the existing methods in accurately annotating cell types and achieves a higher performance overall in both cell type prediction tasks.

### Joint learning and annotation in diverse single-cell multiome datasets

Hydra’s capabilities also extend to joint learning and automated annotation of single-cell multiome datasets, and can simultaneously handle three different single-cell modalities, including transcriptomic, chromatin accessibility, and protein expression data (6, 7, 27–29). To evaluate Hydra’s performance in this context, we utilized datasets generated from seven single-cell multiome technologies across various tissues such as PBMC, Brain, Skin, Kidney, and Embryo. We compared Hydra against three recent methods also developed for handling single-cell multiome data, including MOFA+ (30), UMINT (31), and scMoMaT (32). All three methods provide embeddings and require additional classifiers to assign cell types in the query datasets. UMINT provides a simple neural network classifier similar to the one used by Hydra. We implemented a classifier with a similar architecture to train on the embeddings obtained from MOFA+ and scMoMaT.

For the intra-dataset cell type identification task, we performed five-time repeated random subsampling validation on datasets obtained from four different single-cell multiome technologies, mainly SHARE-seq, SNARE-seq, sciCAR, and CITE-seq (Table 1). Across all datasets, we observed that Hydra achieved higher performance in predicting cell types compared to other methods (Fig. 4a-d, Supplementary 4a-d). Similar to single-cell transcriptomic cell type prediction, other methods performed poorly in annotating cell types with smaller populations. Further, we conducted inter-dataset benchmarking across different batches involving 26 train-test splits (Table 1). This evaluation included datasets generated from simultaneous profiling of two modalities - RNA+ADT (CITE-seq) and RNA+ATAC (10X Multiome) as well as all three modalities - RNA+ADT+ATAC (TEA-seq). Again, Hydra achieved significantly higher performance in predicting cell types across most datasets, with an overall accuracy of 74%, while the second-best performing method, UMINT, achieved an overall accuracy of 60% (Fig. 4e). The superior performance of Hydra across both single-cell unimodal and multiome datasets can be attributed to its ability to effectively identify rare cell populations, which remain a key challenge for existing approaches.

**Fig. 4.**
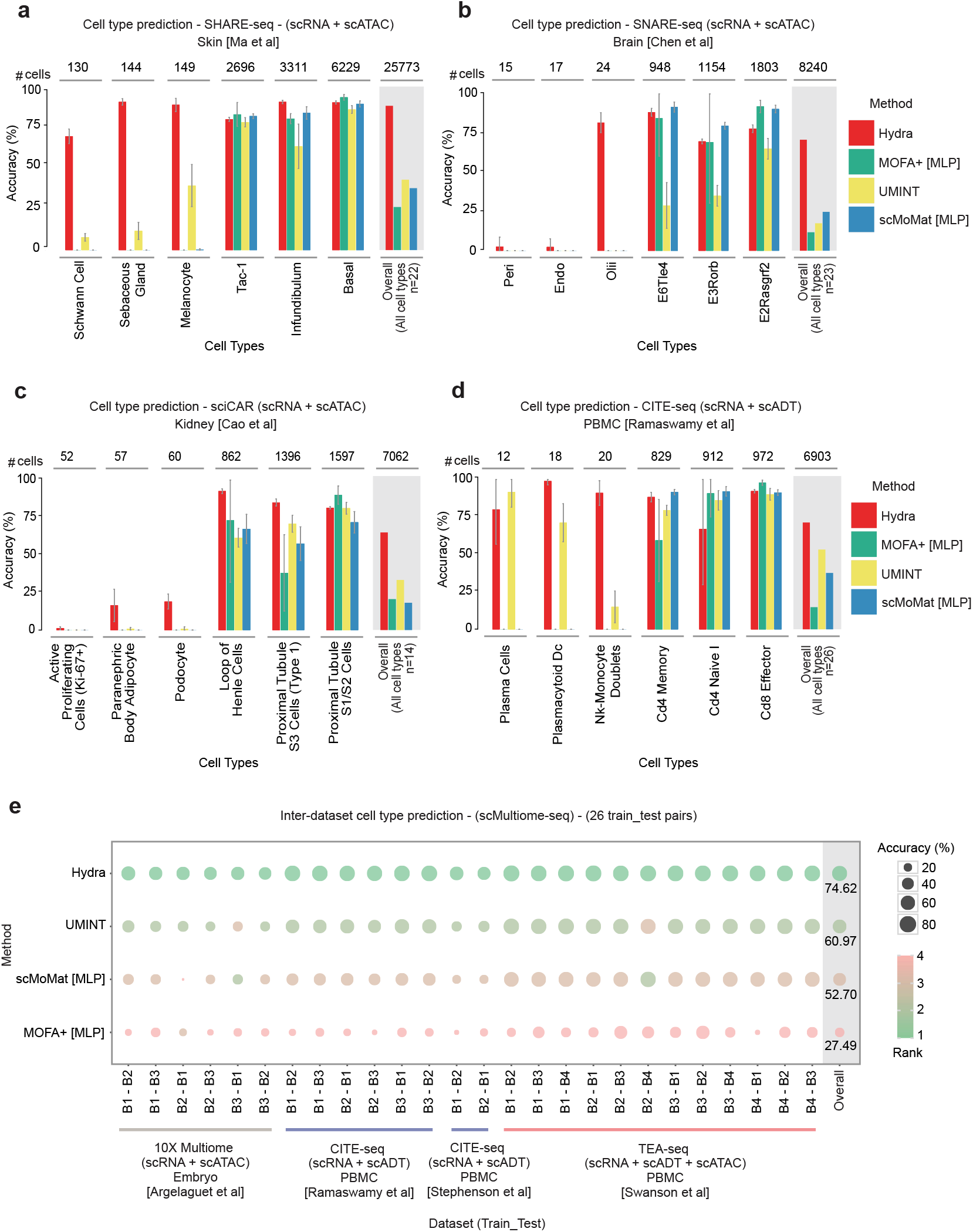
Evaluation of cell type prediction performance of Hydra using single-cell multiome datasets. (a) Bar plot with error bars illustrating the 5-fold intra-dataset cell type prediction performance of Hydra against existing methods using the SHARE-seq Skin dataset. This dataset profiles RNA and ATAC modalities and comprises approximately 25,000 cells across 22 cell types. Prediction accuracy for the three smallest and three largest cell types is displayed, followed by the overall performance across all 22 cell types in the dataset (highlighted with a gray background). (b) Five-time repeated random subsampling intra-dataset cell type prediction accuracy of Hydra against existing methods using the SNARE-seq Brain dataset, which profiles RNA and ATAC modalities and includes approximately 8,000 cells. The overall prediction performance across all 23 cell types is indicated with a gray background. (c) Comparison of five-time repeated random subsampling intra-dataset cell type prediction accuracy of Hydra against other methods using the sciCAR-seq Kidney dataset, which profiles both RNA and ATAC modalities and comprises approximately 7,000 cells. Overall performance across all 14 cell types in the dataset is highlighted with a gray background. (d) Five-time repeated random subsampling intra-dataset cell type prediction performance using the CITE-seq PBMC dataset which profiles RNA and ADT modalities and includes approximately 6,000 cells. The overall performance across all 26 cell types in the dataset is highlighted with a gray background. (e) Visualization of the inter-dataset cell type prediction performance of various methods across four distinct single-cell multiome studies and 26 train-test splits. The bubble plot ranks methods based on their mean accuracy across all datasets.

### Hydra robustly maps cellular subtypes in Alzheimer’s disease from single-cell multiome data

In complex conditions, such as Alzheimer’s disease (AD), significant changes in molecular profiles can often obscure cell identities by altering genetic and epigenetic landscapes. Such changes can make it challenging to accurately identify and distinguish cell types, especially when relying on markers established from a healthy reference. This challenge is particularly acute in the brain, where many neuronal and glial subpopulations share highly similar molecular signatures. Here, we aimed to evaluate Hydra’s robustness in learning from a healthy reference dataset and predicting cellular subtypes across conditions, particularly early- and late-stage Alzheimer’s disease cohorts. To achieve this, we utilized our previously published single-cell multiome dataset that profiled the transcriptome and epigenome of the medial frontal cortex (MFC) region of the brain (33). First, we used an intra-dataset cell type identification approach described earlier using a healthy MFC brain dataset consisting of 27 distinct cell populations, and compared the prediction performance of Hydra against different single-cell multiome methods. We found that Hydra outperformed other methods in predicting both major and minor cell types of the brain with an overall accuracy of approximately 86%. While other methods performed reasonably well in identifying certain major cell types, their performance decreased as the size of the different cell populations decreased (Fig. 5a).

**Fig. 5.**
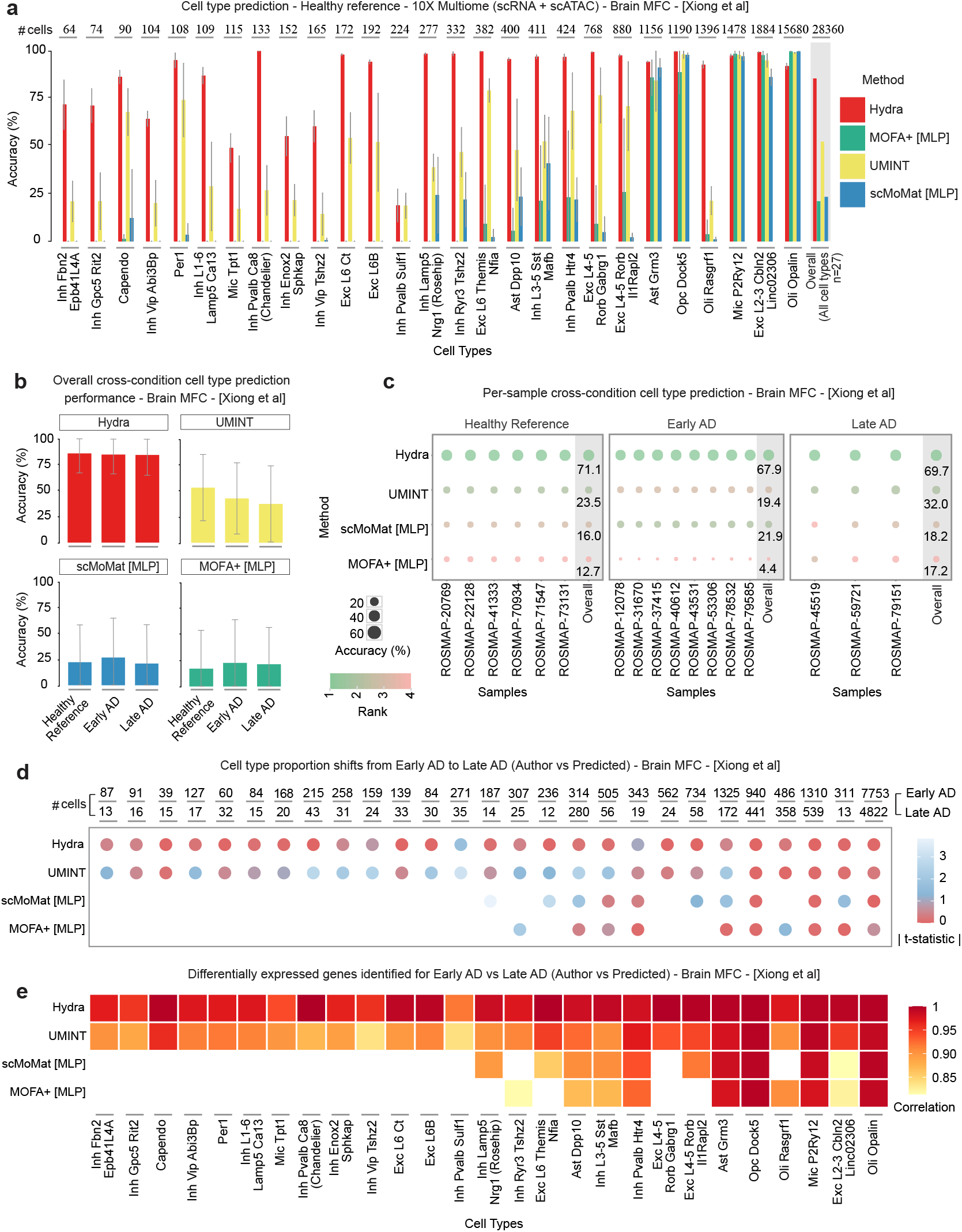
Mapping cellular subtypes of brain medial frontal cortex (MFC) in Alzheimer’s disease. (a) Bar plot with error bars illustrating the 5-fold intra-dataset cell type prediction performance of Hydra against different single-cell multiome methods using healthy medial frontal cortex (MFC) brain dataset. This dataset profiles RNA and ATAC modalities and comprises approximately 28,000 cells across 27 cell types. Prediction accuracy is displayed for all cell types, followed by the overall performance across all cell types in the dataset (highlighted with a gray background). Cell types are arranged in the order of increasing sample count from left to right. (b) Faceted bar plots showing overall cell type prediction performance of different single-cell multiome methods across conditions trained on a healthy reference dataset, stratified by original cell type proportions. Error bars represent the variability in prediction accuracy across individual cell types. (c) Bubble plot illustrating per-sample cross-condition identification of cell types using different single-cell multiome methods trained on healthy samples. The plot is faceted by conditions namely - healthy reference comprising held out healthy samples, early- and late-stage Alzheimer’s disease. Methods are ranked based on overall performance across all conditions. (d) Bubble plot showing absolute t-statistics quantifying cell type proportion changes from early to late stages of Alzheimer’s disease. Each circle represents the magnitude of proportion shift differences between author label and predicted label for each cell type. (e) Heatmap illustrating Spearman correlation coefficients between differential gene expression t-statistics derived from autho9r annotations and method predictions for early versus late AD.

To further evaluate the robustness of methods, we conducted a cross-condition cell type prediction task. Essentially, we trained each method on a subset of the healthy reference dataset stratified by original cell type proportions and used the trained models to predict cell types in held-out healthy reference, early- and late-stage Alzheimer’s disease datasets. Remarkably, Hydra accurately transferred annotations from the healthy reference and identified most cell types in early- and late-stage Alzheimer’s disease, with an overall accuracy of approximately 85% and 84%, respectively. In contrast, the performance of the second best-performing method, UMINT, decreased in early- and late-stages of Alzheimer’s disease. scMoMat and MOFA+ demonstrated consistent but much lower performance across all conditions, similar to that observed in the intra-dataset prediction task above (Fig. 5b, Supplementary 5a). Next, to further evaluate the generalizability across disease-specific sample variations and to demonstrate cross-condition cell-type prediction on unseen samples, we withheld a subset of healthy samples and trained the methods on the remaining healthy cohort. We then applied the trained models to predict cell types in held-out healthy samples as well as in the early- and late-stage of Alzheimer’s disease. We observed that performance of Hydra decreased slightly but still achieved a considerably high cell type prediction accuracy across all conditions in comparison to other methods (Fig. 5c). These results show Hydra’s ability to reliably identify cell type markers from healthy samples and demonstrate its robustness to molecular-level shifts occurring in disease conditions.

Finally, to demonstrate that the predicted cell types capture biologically meaningful changes characteristic of Alzheimer’s disease progression. To answer this, we performed two complementary analyses comparing cell type proportion shifts and differential gene expression patterns between early- and late-stages of AD. First, we quantified cell type proportion changes from early to late AD using sample-level t-statistics and compared the magnitude of changes captured by author annotations with individual method predictions across all cell types. Hydra demonstrated consistent performance in capturing proportion shifts across the majority of cell types, with UMINT showing moderate performance as the second-best method. In contrast, scMoMat and MOFA+ showed substantial gaps, particularly failing to predict several low-abundance cell types, highlighting their limited ability to transfer annotations to rare but potentially important cellular subtypes in disease contexts (Fig. 5d). Subsequently, we assessed whether predicted cell types preserve disease-relevant transcriptional signatures by performing pseudobulk differentially expressed gene (DEG) analysis between early- and late-stages of AD. We calculated Spearman correlations between DEG t-statistics derived from author annotations and those from individual method predictions for each cell type. We found that Hydra outperformed other methods in maintaining strong correlations with author DEG signatures in the identified cell types, including rare cell populations, demonstrating its ability to preserve critical disease-relevant molecular signatures (Fig. 5e).

## Discussion

In this paper, we introduced Hydra as an interpretable deep generative ensemble learning framework designed to automate the process of identifying cell type-specific features and predicting cell types in both unimodal and multimodal single-cell omics datasets. By integrating multiple variational autoencoders, Hydra generates new samples from learned probability distributions to efficiently mitigate class imbalance and ensure that all cell types, including rare/minor cell populations are adequately represented. This approach preserves essential biological signals during model training and enables Hydra to robustly capture cellular markers, especially for rare cell types.

Another key advantage of Hydra lies in addressing the important challenge of interpretability observed with traditional deep learning tools. By employing an Integrated Gradients attribution approach, the feature ranking module ensures model interpretability and reveals the genes that collectively dictate cellular identity. We show that Hydra consistently offers superior performance compared to existing methods across 21 datasets, covering a wide range of single-cell omics modalities and tissues. Furthermore, our evaluation strategy emphasizes predictive performance for each individual cell type, which is particularly important for identifying minor cell populations often overlooked by more generalized global metrics. Finally, we demonstrate Hydra’s ability to facilitate robust reference-to-query cell type mapping in both normal and disease contexts, including independent inter-dataset tasks across different sequencing protocols.

As single-cell multiomics technologies become increasingly accessible, we anticipate that Hydra will serve as a valuable tool to harness such rich, multilayered information and shed light on underlying cellular heterogeneity.

## Methods

### Data Processing and Normalization

The input for Hydra can include both unimodal and multimodal single-cell data. We start with a count matrix *M* of dimensions *n × m*, where *n* denotes the number of cells and *m* denotes the number of features. To reduce noise and retain biologically relevant features, we filter out features *f* if the proportion of cells for which *f*_*i*_ has a zero count is greater than or equal to 99%. Feature *f*_*i*_ is retained if

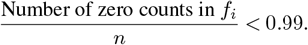

After filtering, the count matrix *M* is logarithmically transformed. Subsequently, the matrix is scaled and transformed into a PyTorch tensor for model input.

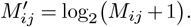

### Feature ranking module

#### Model architecture

To capture the underlying heterogeneity in the single-cell data, we employ a variational autoencoder architecture with multiple classification heads in the initial training phase of the feature ranking module. For unimodal single-cell data such as transcriptomic and chromatin accessibility, the encoder and decoder architecture is consistent with a latent space *z* of 100 neurons. In the case of single-cell protein expression, we use fewer neurons in the hidden layers due to lower feature dimensionality. The single-cell multiome data adopts a similar architecture to unimodal data, but modality-specific encoder layers process each modality independently before concatenating and passing them through a shared latent space *z* of 100 neurons. This joint learning allows Hydra to capture information embedded in different single-cell modalities.

The input feature matrix *x* is first projected through a linear layer, followed by Mish activation (Misra) and batch normalization (Ioffe and Szegedy), with a dropout rate of 0.2. This is followed by a second linear transformation to reduce the dimensionality to the latent space *z*. The reparameterization trick (Kingma and Welling) is applied to sample from *z* during optimization.

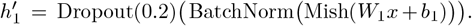

For single-cell multiome data, modality-specific encoders process each input independently. Let *x*_1_ and *x*_2_ represent the input feature matrices from two modalities.

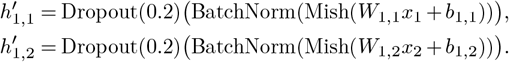

The encoded features from both modalities are concatenated into a combined latent space, which is further processed through a fully connected layer to create the shared latent variable *z* for joint learning. In the decoder, the latent variable *z* is mapped back to the original feature space.

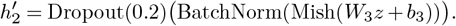

The shared latent variable z is then passed through modality-specific decoders, where each modality undergoes similar transformations. The latent space *z* is also fed into multiple classification heads. The number of classification heads *C*_*m*_ is user-configurable, with a default of 25 set empirically.

#### Initial training phase

During the initial training phase, we employ a dynamic sampling strategy to handle class imbalance in input single-cell data. This approach progressively adjusts the probability of selecting samples from each class as training progresses. Specifically, at each epoch *e*, we compute a weighting factor for each class based on its sample count *n*_*c*_ and the current epoch number. If *E* is the total number of training epochs, the sampling weight *w*_*c*_ for class *c* is calculated as:

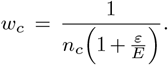

Early in training, when e is small, the weights are inversely proportional to the class counts *n*_*c*_, and higher preference is given to samples from minority classes. As training progresses, the denominator increases, gradually reducing the weights for all classes. Since minority classes have smaller *n*_*c*_, their sampling probabilities decrease more slowly compared to those of majority classes. The weights are normalized, and samples are drawn according to these probabilities in each iteration.

The model optimizes a loss function that takes into account both the reconstruction and classification tasks to capture the underlying heterogeneity in the single-cell data. The reconstruction loss *L*_recon_ consists of the mean squared error between the input *x* and the reconstructed data 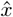, and the Kullback–Leibler divergence L_KL_ for the variational component:

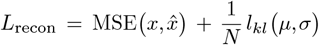

Where *N* is the number of features, *µ* and *σ* are the mean and standard deviation from the encoder’s output. For the classification task, the model employs multiple classification heads *C*_*m*_that operate on the latent space *z*. Predictions from all classification heads are averaged to obtain the final class 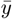. The classification loss L_class_ is then computed using a label-smoothed cross entropy function (34) between the averaged prediction *ŷ* and the true class labels *y*:

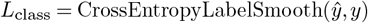

Finally, to optimize the training time, we also implement an early stopping approach with a patience of 10 epochs.

#### Refinement and feature ranking

During the refinement stage, we generate a balanced dataset from the original input single-cell data. We do this by categorizing cell types into major and minor groups based on the median sample size across all cell types. To create a balanced dataset, we randomly down-sample the major cell types and generate additional samples for minor cell types using the original trained model. We then initialize a new model for refinement by inheriting the weights of the original model’s encoder, decoder, and a specific classification head *C*_*m*_. Specifically, we create a new model for each classification head in the original trained model. For each initialized model, we generate a new balanced dataset following the above procedure. The refined models are then trained on their respective balanced datasets for a varying number of epochs. After refining all the models, we apply a post-hoc feature attribution technique, Integrated Gradients (IG), to compute importance scores for each feature in each cell type. To improve computational efficiency, we adopt a batch attribution approach on all samples belonging to a given cell type. For a given cell type *c* and model *m*, the feature importance scores *α*_*m*_ are computed by summing the absolute values of the Integrated Gradients attributions over all samples of cell type *c*:

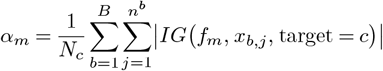

Where *N*_*c*_ is the total number of samples of cell type *c, B* is the total number of batches, *n*^*b*^ is the number of samples in batch *b* such that

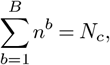

*x*_*b,j*_ is the *j*th sample in batch *b*, and *f*_*m*_ is the model comprising of encoder and classification head *C*_*m*_ from the refined model *m*. The features are then ranked considering the biological directionality. Finally, we obtain a consensus cell type-specific feature ranking by averaging the importance scores across all refined models.

### Automated cell type annotation module

We train an ensemble of simple neural network classifiers to assign cell types in the query single-cell dataset. Each neural network classifier consists of an input layer, a hidden layer with the Mish activation function

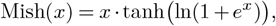

and an output layer. The forward pass through each classifier can be defined as:

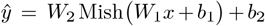

Where *x* is the input feature vector, *ŷ* is the output logits corresponding to the cell type classes, *W*_1_ and *b*_1_ are the weights and biases of the hidden layer, and *W*_2_ and *b*_2_ are the weights and biases of the output layer. The ensemble size corresponds to the number of refined models from the feature ranking module. To train the classifiers, we generate balanced datasets using the refined models similar to the model refinement step. The dimensions of these datasets are reduced using the union of the top 100 ranked features from all cell types, as determined by the feature ranking module. Each neural network classifier is then trained on its respective balanced dataset for five epochs. After training, the ensemble of classifiers is used to map cell types in the query single-cell dataset. Predictions from all classifiers are aggregated by averaging their prediction probabilities to obtain the final cell type assignments.

### Running existing methods

#### Statistical tests

We employed three statistical methods, including Welch’s t-test, Wilcoxon rank-sum test, and Limma-Voom. As a preprocessing step, genes expressed in less than 1% of cells were filtered out. For each cell type, we performed a one-vs-all analysis by comparing the expression levels of genes in the cell type of interest against all other cell types combined. For the t-test and Wilcoxon rank-sum test, raw count matrices were log-transformed and scaled. The test statistics were computed, and genes were ranked considering the biological direction of expression changes. For the Limma-Voom analysis, we used the voom function to model the count data, followed by linear modeling and empirical Bayes moderation using the lmFit and eBayes functions from the Limma package. Cell type-specific genes were ranked based on moderated t-statistics, considering the biological direction.

#### SingleCellNet

SingleCellNet (SCN) was implemented for scRNA-seq cell type identification evaluation following the official documentation (https://pysinglecellnet.readthedocs.io/en/latest/). Genes expressed in less than 1% of cells were filtered out, and raw count matrices were converted into AnnData objects for SCN input. The model was trained using the scn_train function with the following parameters – ‘nTopGenes’ = 200, ‘nTopGenePairs’ = 200, ‘nRand’ = 100, ‘nTrees’ = 1000 and ‘propOther’ = 0.1. Cell types in the query dataset were then predicted using the scn_classify function.

#### scClassify

scClassify was implemented for scRNA-seq cell type identification evaluation, following the official documentation (https://sydneybiox.github.io/scClassify/). Genes expressed in less than 1% of cells were filtered out, and raw count matrices were log-transformed and scaled. Model training and cell type prediction on the query dataset was performed using the scClassify function with the following parameters - ‘tree’ = “HOPACH”, ‘algorithm’ = “WKNN”, ‘selectFeatures’ = c(“limma”), and ‘similarity’ = c(“pearson”, “cosine”).

#### scPred

scPred was implemented for scRNA-seq cell type identification evaluation, following the official documentation (https://powellgenomicslab.github.io/scPred/). Genes expressed in less than 1% of cells were filtered out, and raw count matrices were preprocessed using Seurat’s (35) NormalizeData, ScaleData, and FindVariableFeatures functions. The scPred input object was then created using the getFeatureSpace function of scPred. Model training and cell type prediction on the query dataset was performed using the trainModel and scPredict functions, respectively.

#### scVI

scVI was implemented for scRNA-seq cell type identification evaluation, following the official documentation (https://github.com/scverse/scvi-tools). Genes expressed in less than 1% of cells were filtered out, and raw count matrices were converted to AnnData objects. First, feature selection was performed using Scanpy (36). The setup_anndata function of scVI was then used to prepare the input data. The model was initialized using scvi.model.SCVI and trained with the train function. The latent space of the query dataset was obtained using the trained model’s get_latent_representation function, followed by clustering with Scanpy, and the most frequent cell type label for each cluster was determined.

#### UMINT

UMINT was implemented for single-cell multiome cell type identification, following the official documentation (https://github.com/deeplearner87/UMINT). Genes expressed in less than 1% of cells were filtered out, and the raw count matrices were log-transformed and scaled. The model was trained using the CombinedEncoder function with the following parameters – ‘layer_neuron’ = [128, 10], ‘mid_neuron’ = 100, ‘seed’ = 42, ‘lambda_act’ = 0.0001, ‘lambda_weight’ = 0.001, ‘epoch’ = 25, and ‘bs’ = 16. The encodings were extracted using the predict function. By default, UMINT provides a simple neural network classifier for cell type predictions, similar to the one used by Hydra. The neural network classifier was trained on the embeddings with the following parameters – ‘layer_neuron’ = [900, 1500], ‘loss’ = “categorical_crossentropy,” ‘epoch’ = 25, and ‘bs’ = 8.

#### MOFA+

MOFA+ was implemented for single-cell multiome cell type identification, following the official documentation (https://biofam.github.io/MOFA2/tutorials.html). Genes expressed in less than 1% of cells were filtered out, and the raw count matrices were log-transformed and scaled.

The model options were set using the set_model_options function with the following parameters – ‘factors’ = 10, ‘spikeslab_weights’ = True, ‘ard_weights’ = True, and ‘ard_factors’ = True. The training options were set using the set_train_options function with parameters – ‘iter’ = 100, ‘convergence_mode’ = “fast”, ‘dropR2’ = 0.001, and ‘seed’ = 42. The model was then built and run using the build and run functions, respectively. The embeddings obtained were used to train a simple neural network classifier similar to the one used by Hydra for cell type prediction on the query dataset with the following parameters – ‘loss’ = “categorical_crossentropy,” epoch = 25, and ‘bs’ = 8.

#### scMoMat

scMoMaT was implemented for single-cell multiome cell type identification, following the official documentation (https://github.com/PeterZZQ/scMoMaT).

Genes expressed in less than 1% of cells were filtered out, and the raw count matrices were preprocessed using the quantile_norm and quantile_norm_log functions of scMoMaT. The dataset objects were prepared according to the author’s pipeline. The model was trained using the scmomat_model and train_func functions with the following parameters – ‘K’ = 30 and ‘T’ = 4000. The embeddings were extracted using the extract_cell_factors function. These embeddings were used to train a simple neural network classifier similar to the one used by Hydra for cell type prediction on the query dataset with the following parameters – ‘loss’ = “categorical_crossentropy,” ‘epoch’ = 25, and ‘bs’ = 8.

### Benchmarking tasks

#### Feature stability of Hydra model variants

We utilized stratified random sampling based on original cell type proportions to obtain five different subsets from each dataset. Different model variants of Hydra were then applied to each subset to obtain cell type-specific features. To quantify feature stability within a dataset, we computed the Pearson correlation coefficient of importance scores between every pair of subsets. If *S*_*i*_ and *S*_*j*_ are two different subsets from a given dataset, the Pearson correlation coefficient *r*_*ij*_ was computed as:

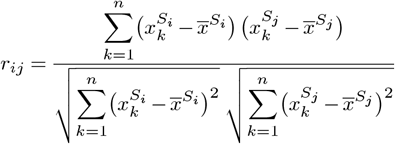

Where 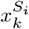 and 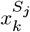 are the importance scores of the *k*th feature in subsets *S*_*i*_ and 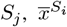 and 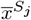 are the mean importance scores in subsets *S*_*i*_ and *S*_*j*_, and *n* is the total number of features evaluated.

#### Feature correlation

We assessed the correlation of top features selected by different feature selection methods using a subsampled dataset with 20 random cell types from the Mouse Cell Atlas. Half of the cell types in the subsampled dataset were considered as major, and the remaining as minor, with a major-to-minor cell imbalance ratio of 100:2. We then applied different feature selection methods to obtain cell type-specific features. The correlation of importance scores was then computed for the top 100 features identified for each cell type. Finally, hierarchical clustering was performed on the correlation matrix to assess similarities in feature importance between cell types.

#### Selecting cell type-specific markers

Similar to feature correlation, we applied feature selection methods on the subsampled dataset from the Mouse Cell Atlas to select markers for the 20 cell types. Next, we randomly sampled 200 cells for each cell type from the original Tabula Muris dataset and examined the expression patterns of the top 100 genes from each cell type. The selection of cell type-specific genes was determined by assessing whether the top genes identified for a cell type of interest exhibited expression specific to that cell type compared to all other cell types.

#### Cell type identification

We divided the cell type identification task into intra-dataset and inter-dataset annotation tasks. For intra-dataset annotation, the dataset was split into train and test sets and evaluated with five-time repeated random subsampling validation, stratified based on original cell type proportions. We then computed the variability in classifying each cell type across different validation subsets. For inter-dataset tasks, cell type identification was evaluated by training on one dataset and testing on an independent dataset, e.g., different batches or studies. The methods were applied to simultaneously predict all cell types in the query dataset. For a given cell type *c*, accuracy was computed as:

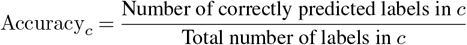

If *N* is the total number of cell types, the overall accuracy was then calculated as the mean of the individual accuracies across all cell types:

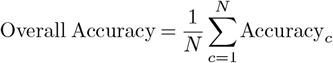

## Data availability

All datasets used in this study are publicly available and a detailed summary of the data is provided in Table 1. The single-cell multiome data, matrices, and metadata profiling Alzheimer’s disease are available at https://compbio.mit.edu/ad_epigenome/.

## Code availability

Hydra framework is implemented as a user-friendly Python PIP package and is available at https://pypi.org/project/hydra-tools/. Documentation for using the tool is available at https://sydneybiox.github.io/Hydra/.

## Acknowledgments

We thank the School of Mathematics and Statistics, The University of Sydney, and the Massachusetts Institute of Technology for providing the high-performance computing resources that have contributed to the research results reported in this paper. This work is supported by scholarships from The University of Sydney and Children’s Medical Research Institute to M.M.W and a USyd Robinson Fellowship to P.Y.. Finally, the authors would like to thank intellectual engagement and feedback from colleagues at the Sydney Precision Data Science Centre, Children’s Medical Research Institute, and MIT Computer Science and Artificial Intelligence Laboratory. They would also like to thank Andy Tran, Lijia Yu, Daniel Kim, Sanghyun Kim, Rajan Shankar, Jackson Zhou, Alex Qin, and Max Wollard for their valuable feedback.

## Author information

Study design - P.Y., E.P., M.M.W., M.K.; Tool development - M.M.W., C.L.; Evaluation and data analyses - M.M.W.; AD data processing – M.M.W., Z.L., Y.W.; Project supervision - P.Y., E.P., M.K.; Manuscript drafting - M.M.W., P.Y., E.P.; All authors read and approved the final version of the manuscript.

---

**Supplementary Figure 1.**
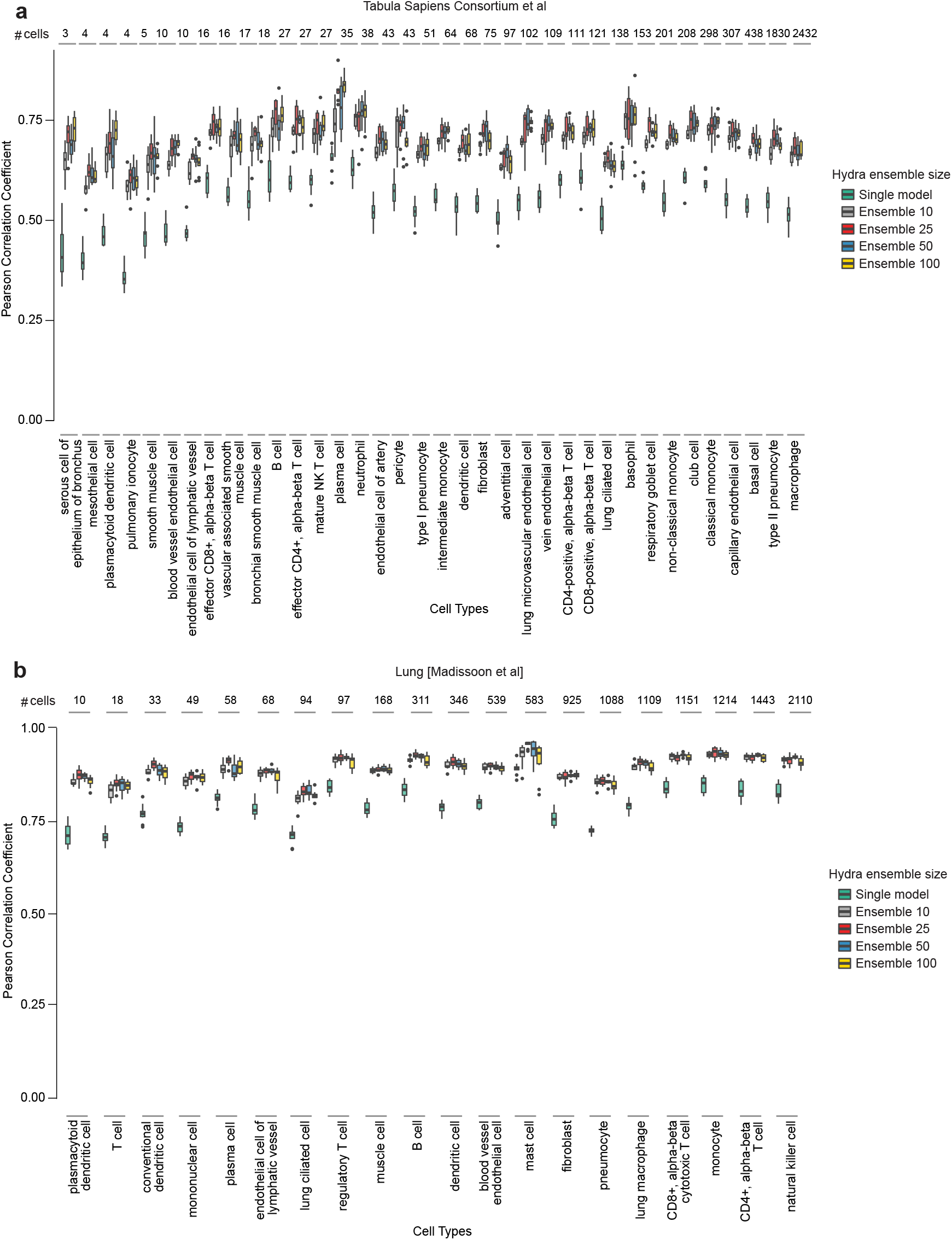
Feature stability of Hydra model variants. (a) The stability of features selected by different model variants of Hydra (n = 1, 10, 25, 50, 100) showing all cell types using the lung dataset (36 cell types), quantified by Pearson correlation coefficients. Cell types are arranged in the order of increasing sample count from left to right. (b) The stability of features selected by different model variants of Hydra (n = 1, 10, 25, 50, 100) showing all cell types using the lung dataset (20 cell types), quantified by Pearson correlation coefficients. Cell types are arranged in the order of increasing sample count from left to right.

**Supplementary Figure 2.**
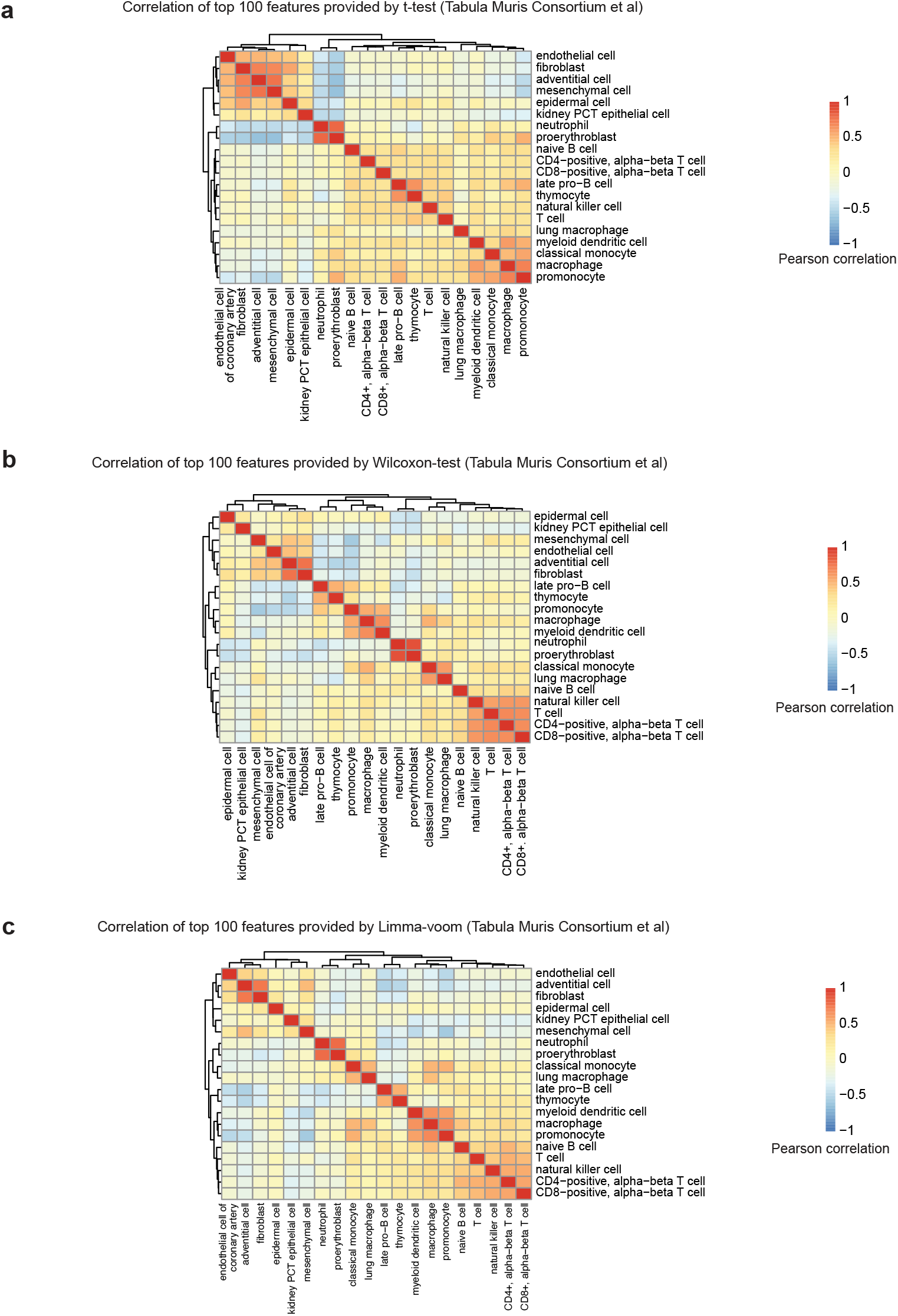
Hierarchical clustering of cell types using the subsampled dataset based on top features selected by statistical methods. Hierarchical clustering of twenty cell types using the top 100 features selected by the t-test (a), Wilcoxon-test (b), and Limma-voom (c) on the subsampled Mouse Cell Atlas. The dataset comprises ten major cell types and ten minor cell types, with an imbalance ratio of major to minor cells of 100:2.

**Supplementary Figure 3.**
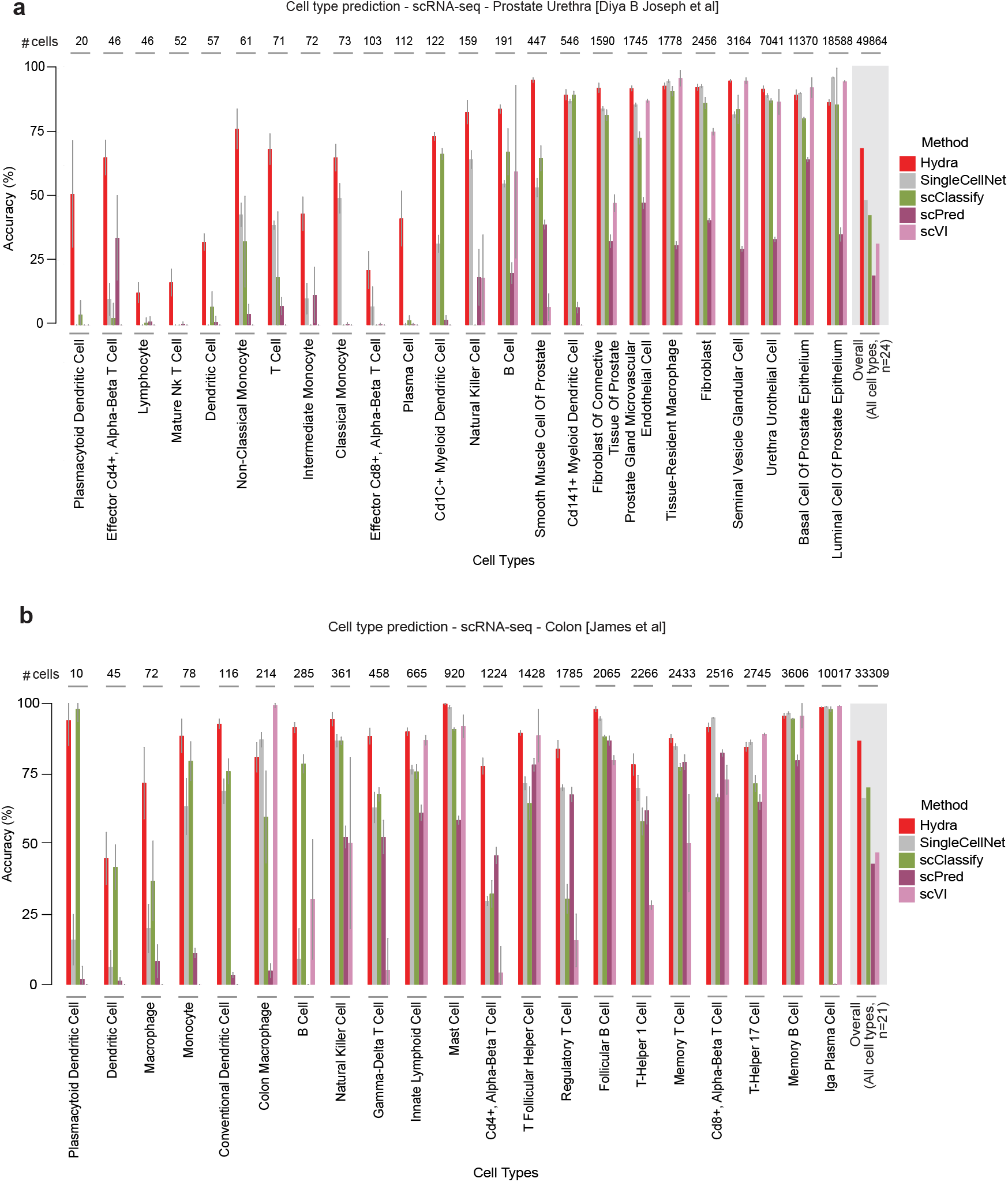
Intra-dataset cell type prediction performance of all cell types in single-cell transcriptomic data. Bar plot with error bars illustrating the five-time repeated random subsampling intra-dataset prediction performance of all cell types using the scRNA-seq Prostate dataset [n=49k cells, 24 cell types] (a) and scRNA-seq Colon dataset [n=33k cells, 21 cell types] (b). Cell types are arranged in the order of increasing sample count from left to right.

**Supplementary Figure 4.**
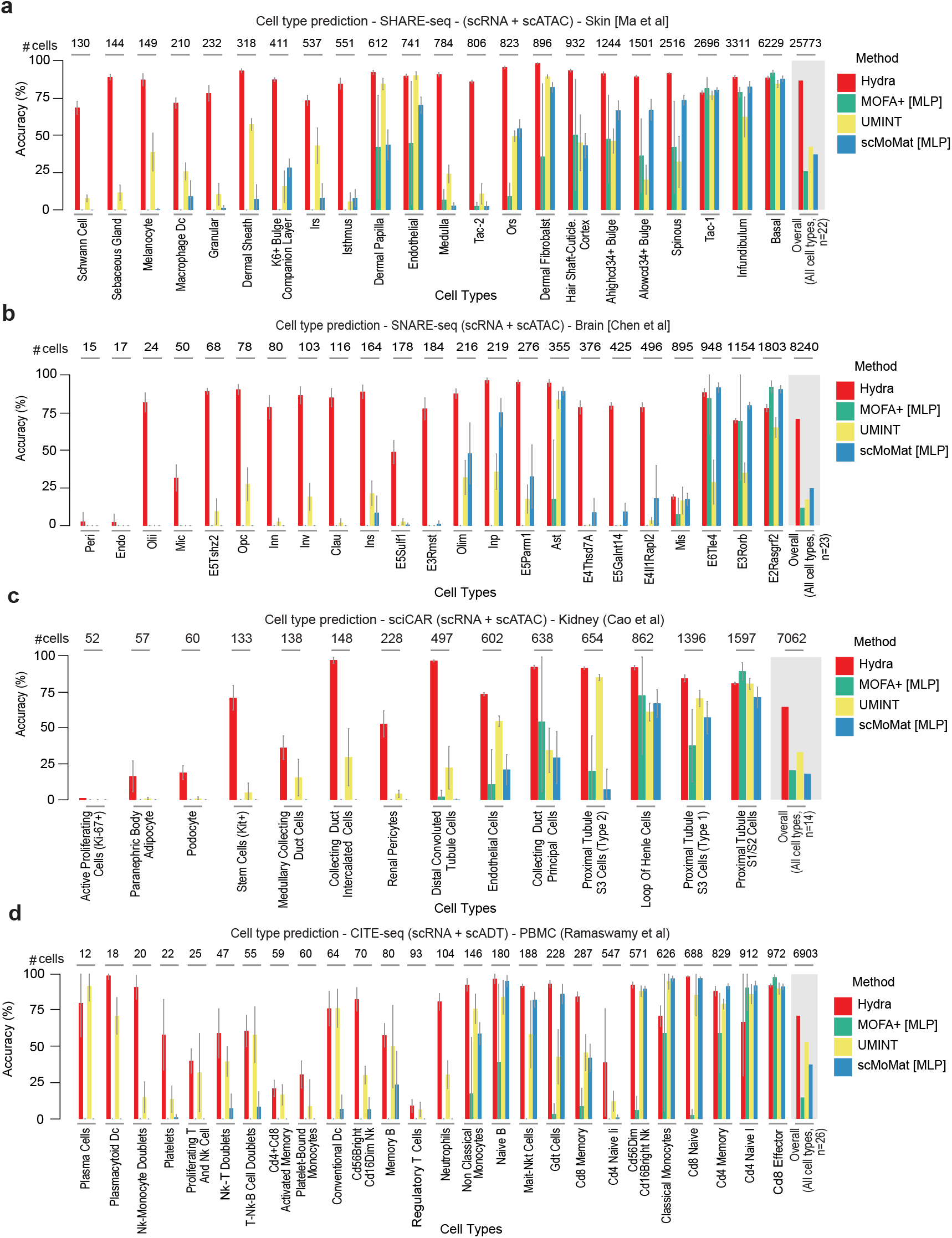
Intra-dataset cell type prediction performance of all cell types in single-cell multiome data. Bar plot with error bars illustrating the five-time repeated random subsampling intra-dataset prediction performance of all cell types using the scMultiome datasets - (a) SHARE-seq Skin [n=25k cells, 22 cell types], (b) SNARE-seq Brain [n=8k cells, 23 cell types], (c) sciCAR Kidney [n=7k cells, 14 cell types] and (d) CITE-seq PBMC [n=6k cells, 26 cell types]. Cell types are arranged in the order of increasing sample count from left to right.

**Supplementary Figure 5.**
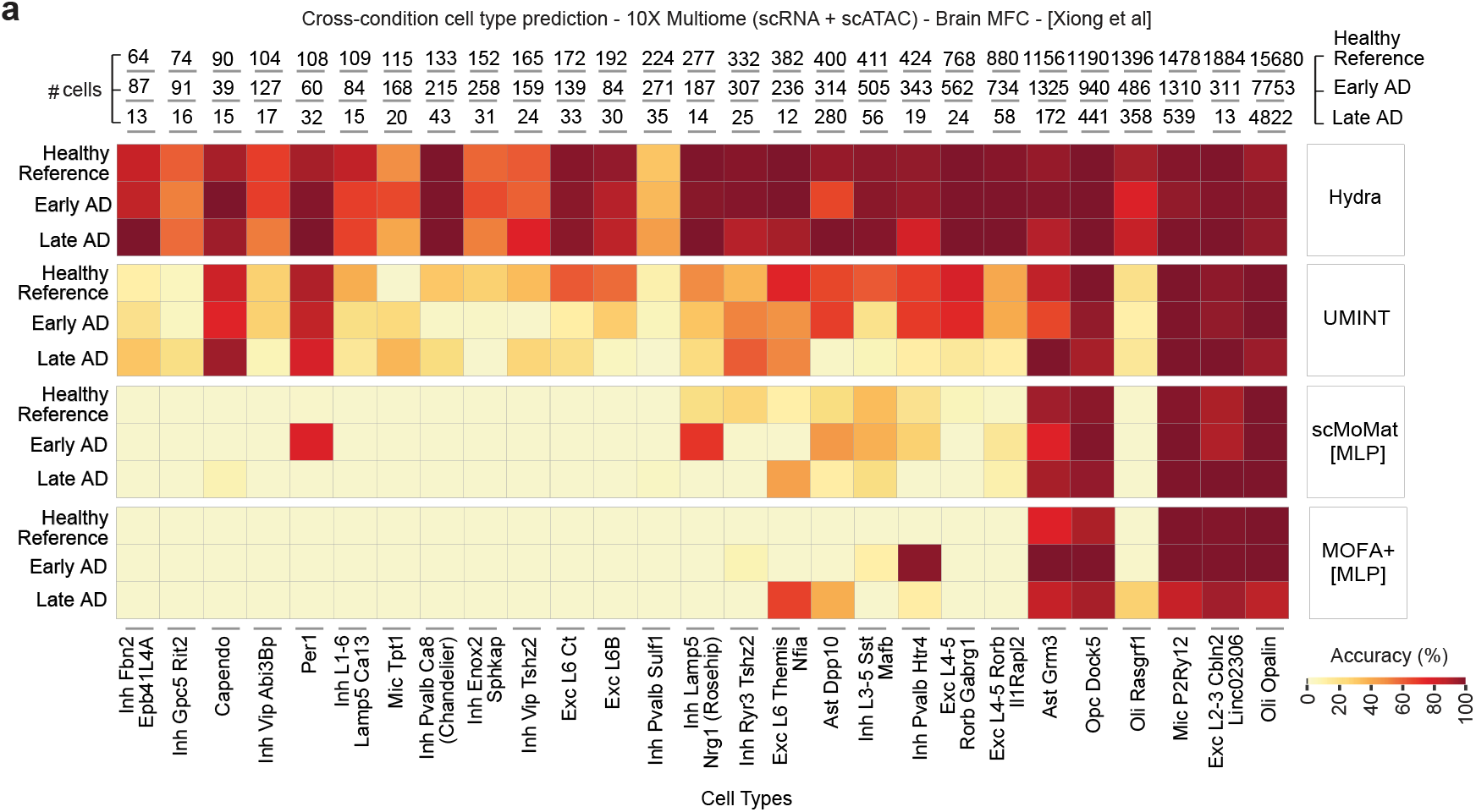
Cross-condition mapping of brain MFC cellular subtypes in Alzheimer’s disease. (a) Heatmaps showing cross-condition cell type prediction performance of different single-cell multiome methods using models trained on healthy reference dataset, stratified by original cell type proportions. Cell types are arranged in the order of increasing sample count from left to right based on healthy reference.

